# The Gut Microbiota composition of Feral and Tamworth Pigs determined using High-Throughput Culturomics and Metagenomics Reveals Compositional Variations When Compared to the Commercial Breeds

**DOI:** 10.1101/738278

**Authors:** Gavin J. Fenske, Sudeep Ghimire, Linto Antony, Jane Christopher-Hennings, Joy Scaria

## Abstract

Bacterial communities in the hindguts of pigs have a profound impact on health and disease. Yet very limited studies have been performed outside intensive swine farms to determine pig gut microbiome composition in natural populations. Feral pigs represent a unique situation where the microbiome structure can be observed outside the realm of modern agriculture. Additionally, Tamworth pigs that freely forage were included to characterize the microbiome structure of this rare breed. In this study, gut microbiome of feral and Tamworth pigs were determined using metagenomics and culturomics. Tamworth pigs are highly dominated by Bacteroidetes primarily composed of the genus *Prevotella* whereas feral samples were more diverse with almost equal proportions of Firmicutes and Bacteroidetes. In total, 46 distinct species were successfully isolated from 1000 colonies selected. The combination of metagenomics and culture techniques facilitated a greater retrieval of annotated genes than either method alone. Furthermore, the naturally raised Tamworth pig microbiome contained more number of antibiotic resistance genes when compared to feral pig microbiome. The single medium based pig microbiota library we report is a resource to better understand pig gut microbial ecology and function by assembling simple to complex microbiota communities in bioreactors or germfree animal models.

## Introduction

The microbiome in the hindgut of mammals has been associated with feed conversion efficiency (Singh, et al. 2014: 145-54), pathogen exclusion (Piewngam, et al. 2018: 532-7), and the production of metabolites that directly influence host signaling pathways (Byndloss, et al. 2017: 570-5). It has become clear in recent years, that the microbiome has a drastic impact on host health. Many current methods to study the swine microbiome, are based upon dietary intervention (Hedegaard, et al. 2016: e0147373, Metzler-Zebeli, et al. 2015: 8489). That is, a dietary substrate is introduced to the animal and an effect on microbiome composition, typically 16s rRNA analysis, is measured. Within the swine industry, there is an upswell of work devoted to increasing feed conversion rate; feed alone accounts for nearly 60% of production costs (Jing, et al. 2015: 11953). While focusing on feed conversion efficiency makes economic sense, the process disregards the biological factors that shaped hindgut evolution, and thus the evolution of the microbiome in pigs. Much in the same way that sampling of traditional hunter-gatherers has provided insight into the microbiome of humans outside the realms of modern dietary practices (Smits, et al. 2017: 802). Feral pigs in the American South may provide a model of the pig microbiome outside the realm of modern agricultural processes.

Currently there are an estimated 6 million feral pigs in the United States (USDA 2018). Feral pigs were first introduced in the early 1500s by Spanish settlers and cause significant ecological damage. It has been shown that feral pigs decrease the amount of plant litter and cover in areas they feed (Siemann, et al. 2009: 546-53). Yet, with the ecological and economic toll feral pigs exert, little study has been conducted to elucidate the structure of their microbiome. Here, we used feral pigs as a case study to compare against Tamworth breed pigs. The Tamworth breed is thought to be descended from the Old English Forest pig and has not been crossed or improved with other breeds since the late 18^th^ Century (British Pig Association n.d.). The breed is not a traditional animal used in high production agriculture, bred instead for its tolerance to cold weather and ability to forage. Given Tamworth’s unique heritage, close relation to an indigenous pig species in the British Isles, and dietary habits closely matching wild pigs, we chose to include them as another model of the pig microbiome outside of the influence of modern agriculture. Additionally, the Tamworth breed is under watch by the Livestock Conservancy, after previously being designated as threatened, and the microbiome composition has yet to be characterized.

Here we attempt to characterize the microbiomes of Tamworth and feral pigs using metagenomic sequencing and high throughput culturomics on direct colon and cecum contents. To date, modern culturomic efforts have been reserved almost exclusively to human fecal samples and we look to extend such methodology to pigs. The culture strategy employs a single medium with various selection screens to shift the taxa retrieved. A single medium isolation strategy will facilitate downstream defined community studies. For example, simple to complex bacterial communities can be assembled in bioreactors to study the mechanisctics of pig gut microbiome succession(Auchtung, et al. 2015: 42). Similiarly, colonization of such defined communities constituted from a well characterized gut microbiome library could reveal how gut bacterial species or combinations impact gut development and immunity(Goodman, et al. 2011: 6252-7). We further characterized the representiative species genomes from our library by whole genome sequencing. Availability of a well characterized strain library with genome information will facilitate future studies to better understand the role of pig gut microbiome in health and disease.

## Materials and Methods

### Sample Collection and Preparation

Permission was granted from purchasers of three Tamworth pigs to obtain colon and cecum samples immediately following slaughter. Small incisions were made into either the colon or cecum with a sterile disposable scalpel. Lumen contents were gently squeezed into sterile 50 mL tubes, mixed with an equal proportion of 40% anaerobic glycerol (final concentration 20% anaerobic glycerol), and immediately snap frozen in liquid nitrogen. For culture preparation, samples were pooled under anaerobic conditions in a vinyl chamber (Coy Labs, USA). Feral samples were kindly provided by boar hunters in Texas, US. A similar procedure was followed where colon and cecum samples were taken immediately following evisceration, mixed with anaerobic glycerol and frozen.

### Metagenomics

DNA was extracted from gut samples using the DNeasy PowerSoil kit (Qiagen, Germany) following the provided kit protocol. After extraction, Microbial DNA was enriched with the NEBNext^®^ Microbiome DNA Enrichment Kit (New England Biolabs, US) to remove host DNA present after DNA extraction. Metagenomic sequencing was conducted on the Illumina MiSeq platform utilizing V2 (250 bp) paired-end sequencing chemistry. Raw sequencing reads were quality controlled using the read-qc module in the software pipeline metaWRAP (Uritskiy, et al. 2018: 158). Briefly, reads are trimmed to PHRED score of > 20 and host reads not removed by enrichment were removed by read-mapping against a reference pig genome (GCF_000003025.6). Resultant reads from read-qc are hereby referred to as high-quality reads. High-quality reads were passed to Kaiju (Menzel, et al. 2016: 11257) for taxonomy annotation against the proGenomes database (http://progenomes.embl.de/, downloaded March 1, 2019). Kaiju was run in default greedy mode and resultant annotation files were parsed in R (R Core Team 2019). Mash (Ondov, et al. 2016: 132) was run to estimate the Jaccard distance between samples. 10,000 sketches were generated for each sample and the sketches were compared using the *dist* function provided in the Mash software.

Antimicrobial resistance (AMR) genes were predicted from metagenomics assemblies. High-quality sequencing reads were assembled into contigs using the assembly module in metaWRAP; metaSPAdes (Nurk, et al. 2017: 824-34) was the chosen to assemble the reads: contigs greater than 1,000 bp were retained. Prodigal (Hyatt, et al. 2010: 119-) was run to predict open reading frames (ORF) using the metagenomic training set. Abricate (Seemann 2018) was then run to annotate the ORF against the NCBI Bacterial Antimicrobial Resistance Reference Gene Database (https://www.ncbi.nlm.nih.gov/bioproject/PRJNA313047, downloaded April 22, 2019).

Contigs were gathered into bins using three methods: MetaBAT2 (Kang, et al. 2019: e27522v1), MaxBin2 (Wu, et al. 2016: 605-7), and CONCOCT (Alneberg, et al. 2014: 1144). Contig bins were kept if the contamination was less than 5% and bin completeness was greater than 85% as determined by CheckM (Parks, et al. 2015: 1043-55). Bins from the three methods used were refined into a coherent bin set using the bin_refinement module in metaWRAP. Refined bins were reassembled with a minimum contig length of 200 bp and the same contamination and completeness parameters as initial bin construction. Metagenomic bin and pure isolate phylogeny was generated using UBCG (Na, et al. 2018: 280-5) to identify and align 92 marker genes. Tree construction was conducted using RAxML (Stamatakis 2014: 1312-3): GTR+G4 nucleotide model. To identify KEGG homologues, ORF were identified in metagenomic assemblies, bins, and culture genomes using Prodigal. The resultant ORF were annotated against the KEGG database using KofamKOALA (Aramaki, et al. 2019: 602110) run locally.

### Culturomics

Colon and cecum samples were pooled respective to feral and Tamworth samples before culture experiments. All culture experiments, including pooling, were conducted under anaerobic conditions inside an anaerobic chamber (Coy Labs, USA). Samples were serially diluted in sterile anaerobic PBS and spread plated onto the media conditions listed in supplemental table 1. Plates were inoculated at 37°C for 48 hours before initial colony selection. 25 colonies were non-selectively sub-cultured from the initial plate to yBHI plates. The procedure was repeated after 72 hours for a total of 50 colonies per media condition. Colonies were primarily identified using MALDI-TOF (Bruker, Germany). MALDI-TOF scores greater than 2.0 were considered a positive species identification. Scores between 1.7 – 2.0 were taken as positive genus identification. Colonies without a positive MALDI-TOF identification were identified by sequencing the 16s rRNA gene. Briefly, DNA was extracted from colonies using the DNeasy Blood and Tissue Kit (Qiagen, Germany) following the manufacturer’s protocol. 16s rRNA sequence was amplified using 27F and 805R primers. The primer sequence is listed in supplemental table 1. Genomes of the selected strains were sequenced on the MiSeq platform utilizing paired-end v3 chemistry (300 bp). Sequencing reads from individual strains were assembled with Unicycler (Wick, et al. 2017: e1005595): minimum contig length of 200 bp. The raw sequencing reads from the culture isolates and metagenomic samples are hosted at NCBI under the BioProject ID PRJNA555322.

## Results

### Tamworth and feral pigs harbor distinct microbiotas

We chose to examine Tamworth breed and feral pigs as their lifestyles differ from traditionally raised agricultural breeds. The Tamworth pigs sampled here were not given any antibiotics or growth promoters and could freely graze. We hypothesized that such raising would cultivate a microbiota that would be different to that of swine raised in intensive hog farms. To begin the investigation, colon and cecum samples were metagenomically sequenced from both breeds. Figure one shows the taxonomic annotation of the metagenomic reads respective to the source of isolation. Contradicting our hypothesis, Feral and Tamworth pigs have inverse Bacteroidetes to Firmicutes compositions. The phylum Bacteroidetes represents nearly 53% of all classified reads in Tamworth pigs compared to 29% in feral pigs. The abundance of Firmicutes in Tamworth samples is lower than feral samples at 15% and 28% respectively. Additionally, nearly 10% more of the feral reads were unclassified compared to Tamworth (37%, 28%) indicating more of the diversity in feral is not yet known in the proGenomes database. Turning to the genus level, the large increase of Bacteroidetes in Tamworth pigs is primarily composed of the genus *Prevotella*, Figure 1 (B) (38%, feral 11%). Remarkably, the genus *Bacteroides* showed almost identical distribution between the feral and Tamworth pigs (7.6% and 7.6% respectively). The increase of Firmicutes in feral samples is due to an increase in several genera such as *Ruminococcus, Clostridium*, and *Eubacterium* corresponding with significantly higher Shannon diversity index values compared to Tamworth (p = 0.0024, Wilcoxon rank-sum test). Full phylum and genus annotation tables are provided in supplemental table 2.

**Fig 1.**
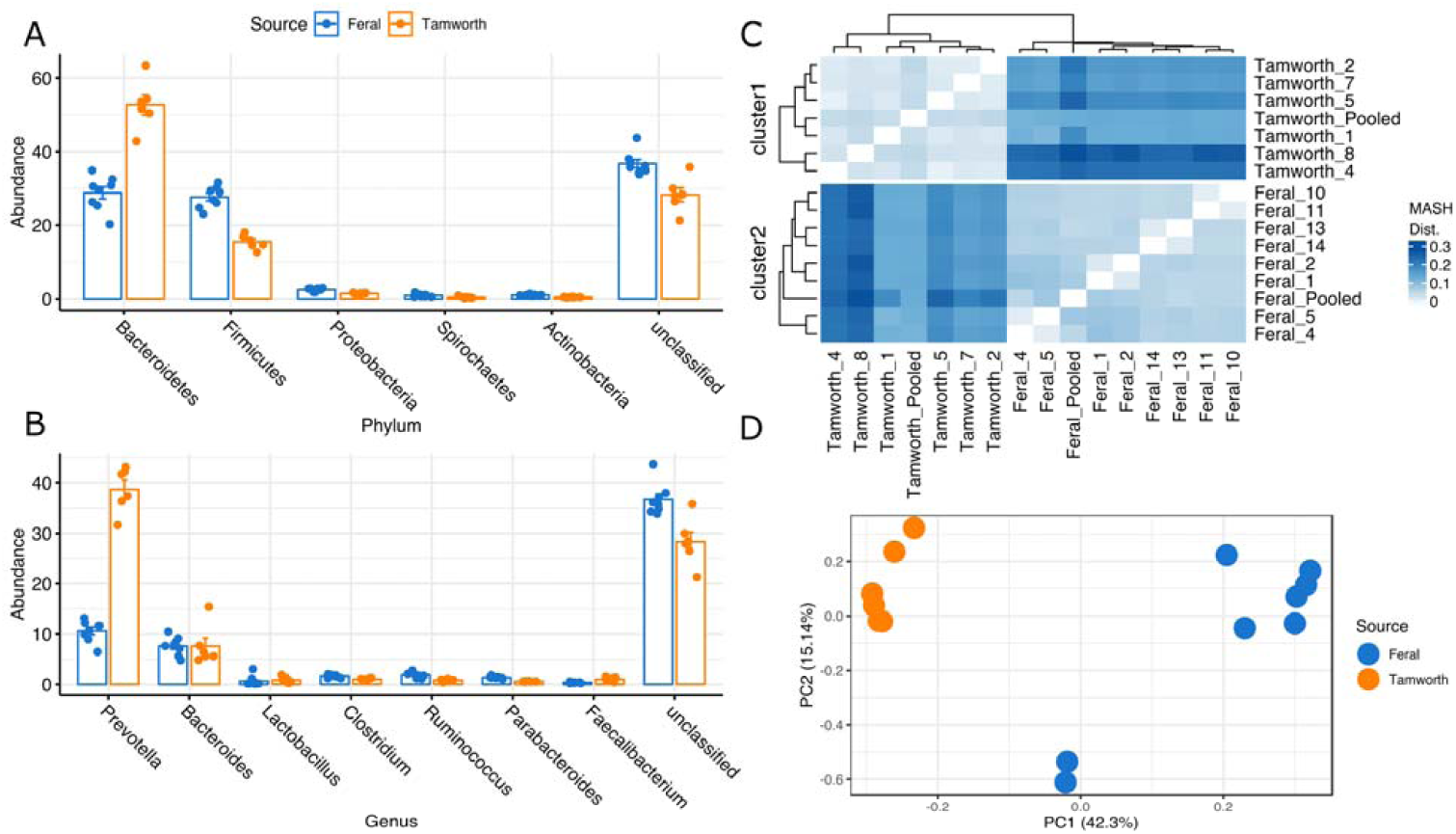
Metagenomic analysis of feral and Tamworth colon and cecum samples. (A)(B) Relative abundance of major phyla and genera annotated from sequencing reads respective to isolation source. (C) Triangle matrix depicting the MASH distance between feral and Tamworth samples. Clusters 1 and 2 are defined by kmeans clustering.

To better understand the distance within a sample source, Tamworth vs. Tamworth, and the difference between sources, Tamworth vs. feral, two clustering methods were employed. First, Mash (Ondov, et al. 2016: 132) was used to sketch the reads sets and compile a distance matrix (Figure 1C). Within the matrix, both Kmeans clustering and hierarchical clustering (average-linkage) separate the samples into Tamworth and feral clades. Mash provides a method to compare metagenomes that is not subject to annotation bias. Principal component analysis (PCA) of the OTU tables was the second method employed. Again, two distinct groups of Tamworth and feral samples are seen in the plot (figure 1D). Interestingly, the Tamworth samples are more homogenous in both the Mash and PCA methods. All pigs were taken from the same farm and this may account for the lower inter-animal microbiome divergence. Thus, the Tamworth and feral pigs examined here harbor distinct microbiotas. Tamworth samples are dominated by the phylum Bacteroidetes, which in turn is largely comprised of *Prevotella*. Feral samples show a much more even distribution of Firmicutes and Bacteroidetes and are more diverse in general.

Stated earlier, Tamworth pigs sampled were not given antibiotics in feed nor given any growth promoters. We hypothesized that the lack of antimicrobial agents would correspond to a relatively low number of AMR homologues in the Tamworth microbiota. Additionally, as feral animals (we presume) do not uptake antimicrobials, their AMR number would be low as well. Confoundingly, Tamworth pigs’ microbiomes contain at least eight AMR homologues. Additionally, all Tamworth samples yield more AMR homologues than feral samples (figure 2). All Tamworth samples contain four putative AMR genes: *cfxA, lnu(AN2), mef(En2), and tet(40).* No common pattern is apparent for Feral samples; *tet(Q)* is found in 5 of 9 feral samples. Thus, in microbiome composition and AMR presence Tamworth pigs do not mirror feral pigs in microbiome composition nor structure. The full result of the antibiotic query is listed in supplemental table 3.

**Fig 2.**
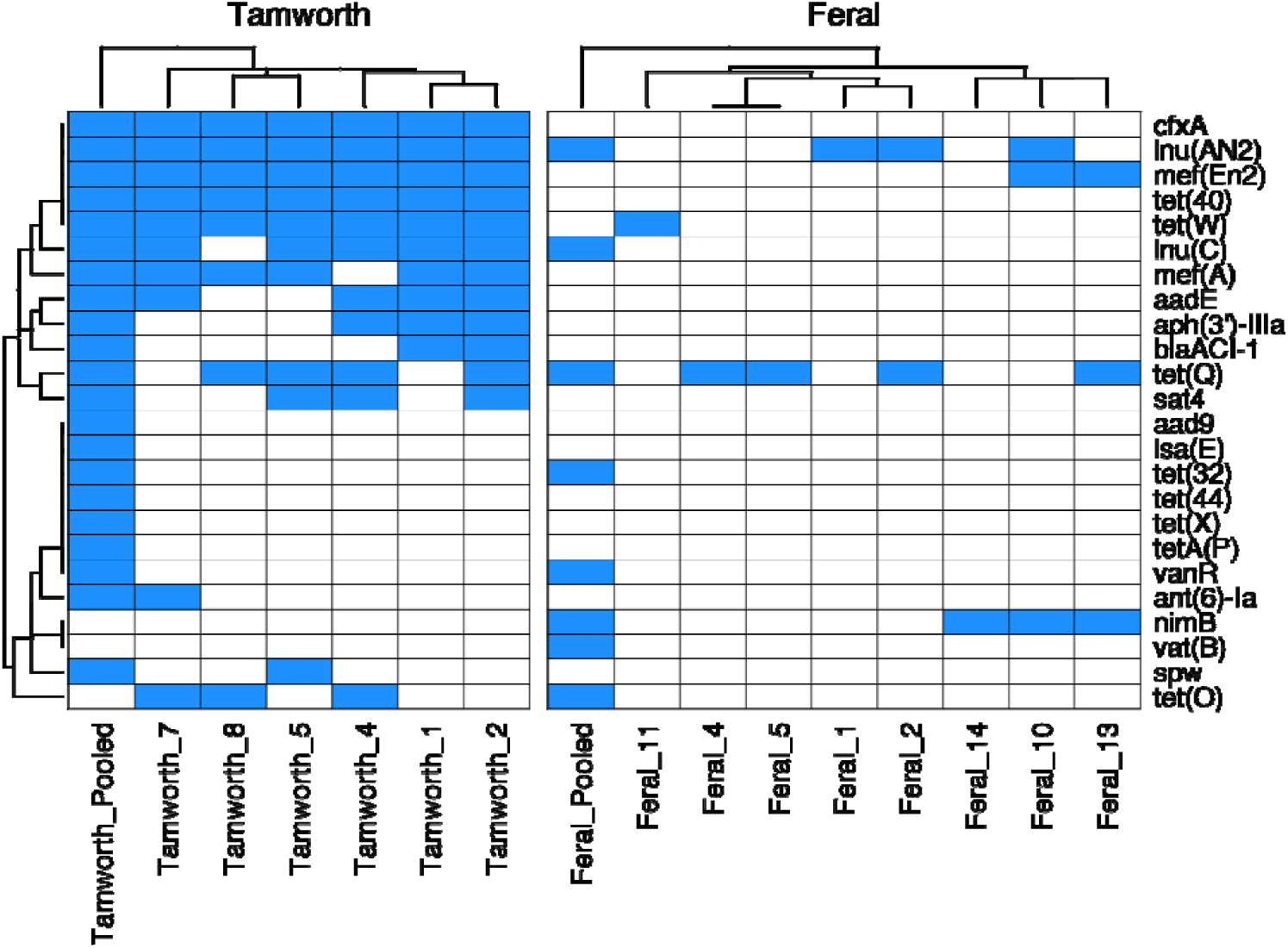
Antimicrobial resistance (AMR) homologues annotated from metagenomic samples. Columns depict individual samples and rows correspond to AMR homologues. Blue color depicts the presence and white color corresponds to absence. AMR homologues were considered present if the coverage value was greater than 90% and a percent homology greater than 70%.

### Selective screens shift plating diversity

High throughput culturomics was the second method employed to sample the two microbiomes. We chose culture sampling, in addition to sequencing methods, as we believed that many low abundance taxa could be retrieved through culture methods that would be lost in metagenomics. Also, the generation of a culture library enables defined community experiments in the future. The culture sampling strategy utilized is as follows: a base medium (yBHI, or close derivatives) had various selective screens (antibiotics, heat, bile, etc.,) applied to it. A finite growing surface is available for colonization and some species will grow more rapidly and subsequently outcompete others. If appropriate selective pressure is applied, we hypothesized that interspecies selection would decrease allowing for taxa not retrieved in plain medium conditions to grow. The approach is similar to one previously used to culture strains from human fecal samples (Rettedal, et al. 2014: 4714). One major difference is said work used multiple media compositions, rather than one as in our study. Ten media conditions were used for both Tamworth and feral samples and are listed in supplemental table 1. 25 colonies were picked at 48- and 72-hours post inoculation, for a total of 50 colonies per condition. In total, 1000 colonies were selected from plates, of which 884 were successfully identified. Selective screens shifted the taxa retrieved (figures 3). Figure 3 depicts the number of isolates per media condition with a bar plot depicting the total number of isolates retrieved. *Lactobacillus sp*. was the most abundant organism retrieved (166 isolates) followed by *Escherichia coli* (86), *Lactobacillus mucosae* (74) and *Streptococcus hyointestinalis* (64). The top ten isolates cultured are listed in table 1. One case of selection completely changing plate diversity compared to plain media is that of heat shock treatment. As expected, many spore forming genera including *Bacillus* and *Clostridium* were only able to grow when the inoculum was heated to kill vegetative cells. The selective screens placed upon yBHI not only shifted the taxa retrieved from each plating condition as shown in Figure 3, but also shifted species richness and evenness (figure 4). The most diverse plating condition (Shannon Index) for both Tamworth and feral samples was obtained from plain yBHI: showing as a log-normal community distribution. Similar log-normal community structures are observed for BSM (Tamworth only), Erythromycin and heat shock treatments. Bile treatments and chlortetracycline exhibited strong selective pressure shown as geometric series in the species-rank abundance plots (figure 4). Most of the taxa retrieved from the bile condition were identified as Proteobacteria, indicating that the dosage of bile (1 g / L) was too high.

**Fig 3.**
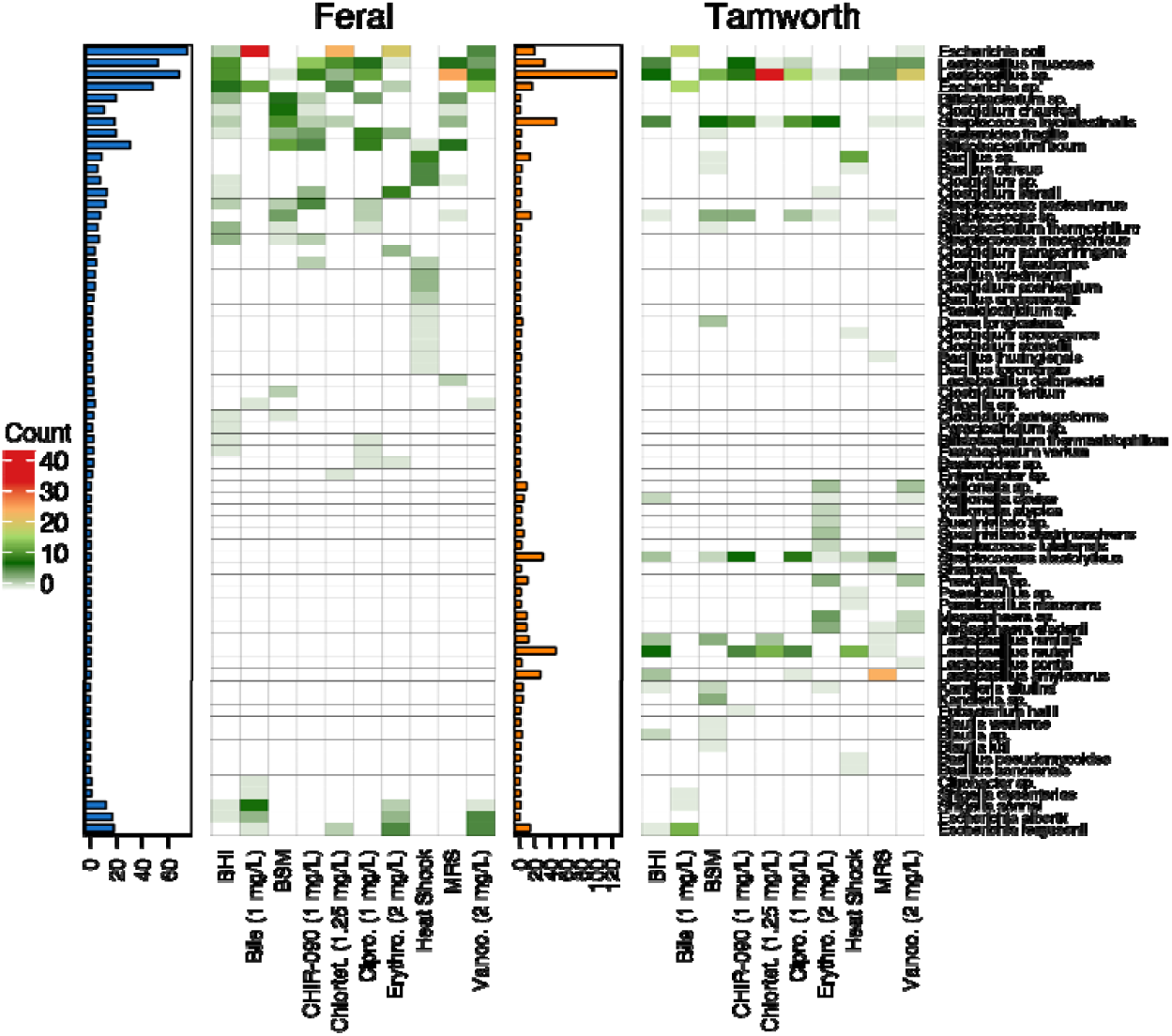
Bacteria isolated from various media conditions. Columns represent individual media conditions and row correspond to bacterial taxa retrieved, cells are colored respective to the number of isolates cultured per media condition. The corresponding bar plot to the left of the matrices shows the total number of isolates retrieved per isolation source.

**Fig 4.**
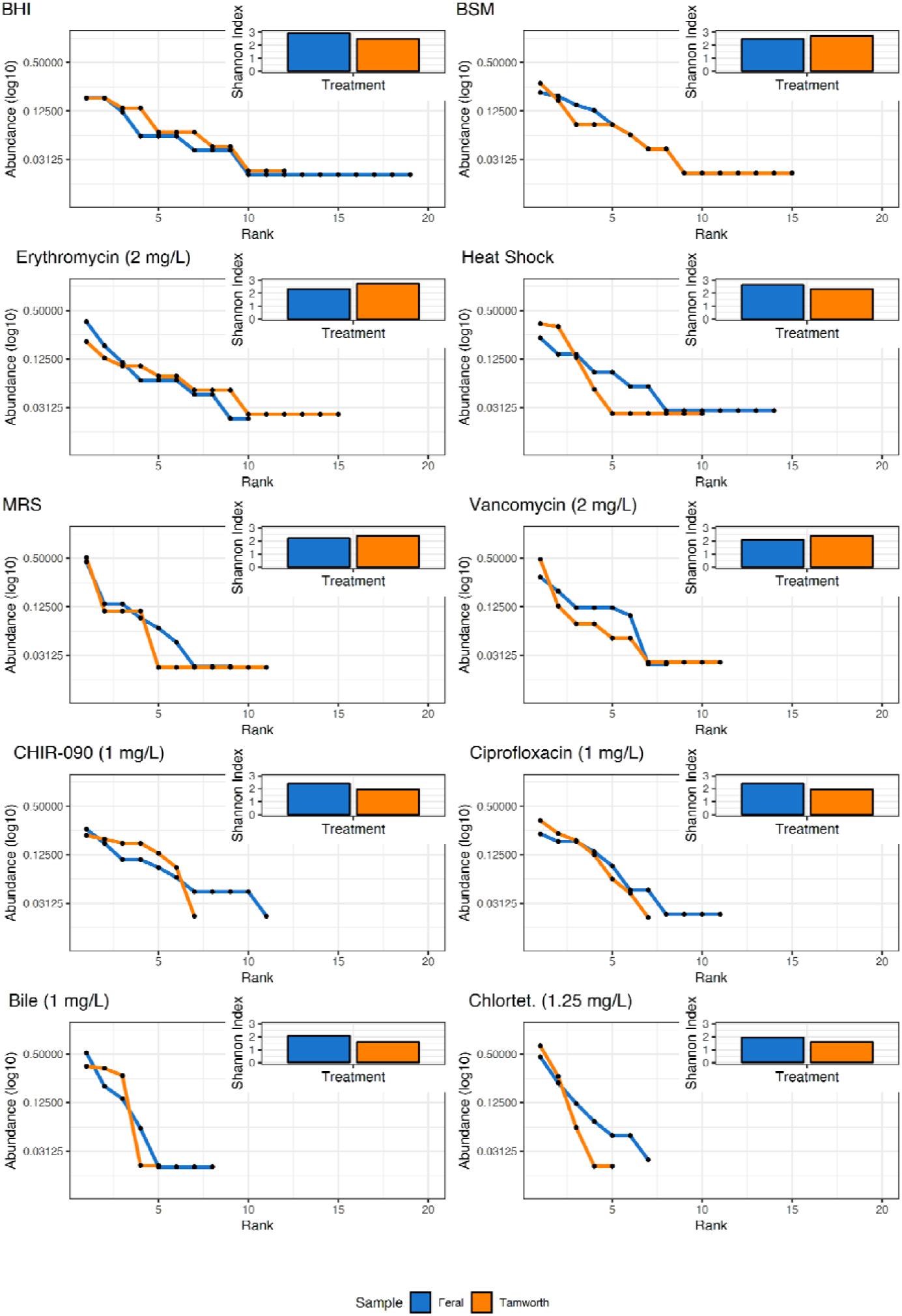
Rank abundance curves of the various media conditions. The community evenness of the various media conditions is shown respective to the isolation source. The inlay plot depicts the Shannon Index respective to the isolation source.

The culture strategy did not recapitulate the community in the inoculum as defined by metagenomics. In both Tamworth and feral samples, a high number of Firmicutes and Proteobacteria were isolated, compared to the metagenomic sampling where Bacteroidetes was the most abundant phylum for both sources. If we disregard the bile conditions, which were dominated by Proteobacteria, yBHI clearly selects for common Firmicutes genera including: *Lactobacillus, Streptococcus*, and *Bacillus.* While the screens were successful in increasing the total number of species retrieved, no condition matched the inoculum in form. *Prevotella* for example, the most abundant genus in Tamworth pigs, was only retrieved seven times from 500 colonies. Taken together, the strategy was successful in gathering many isolates that can grow on a common medium but failed in that the most abundant taxa were not retrieved in proportion to the inoculum.

### Culturing captures genomic information not captured in metagenomics

The sampling strategy employed did not recapitulate the inoculum community. However, one of the main reasons we chose to culture was that we believed rare taxa would provide information that would be loss to metagenomics. To examine this, we sequenced selected isolates and generated 81 high quality metagenomic bins (completeness > 85%, contamination < 5%). The phylogeny of the metagenomic bins and culture genomes was estimated (figure 5). Consistent with read taxonomy, many of the bins constructed from both Tamworth and feral samples were annotated to the phylum Bacteroidetes. The phyla Firmicutes, Proteobacteria and Actinobacteria were comprised almost entirely of isolate genomes. Isolate genomes not only populated clades of the tree missed by metagenomic bins, but provided genes not observed in metagenomic assemblies nor bins (figure 6). Open reading frames (ORF) were predicted from metagenomic assemblies, metagenomic bins, and culture isolate and were annotated against the KEGG database. Figure 6 shows the abundance (natural log) of KEGG homologues respective to the source of the ORF. The full KEGG annotations from the bins, isolates, and metagenomes are provided in supplemental table 4. Metagenomic bins contained less information than the metagenomic assemblies. This is expected as the bins are derived from contigs in the assemblies and not all of the contigs will be gathered into bins. The isolates however provided KEGG homologues that were completely missed through culture-independent methods. Thus, culture and culture-independent methods can augment a microbiota analysis providing information that the other method cannot capture.

**Fig 5.**
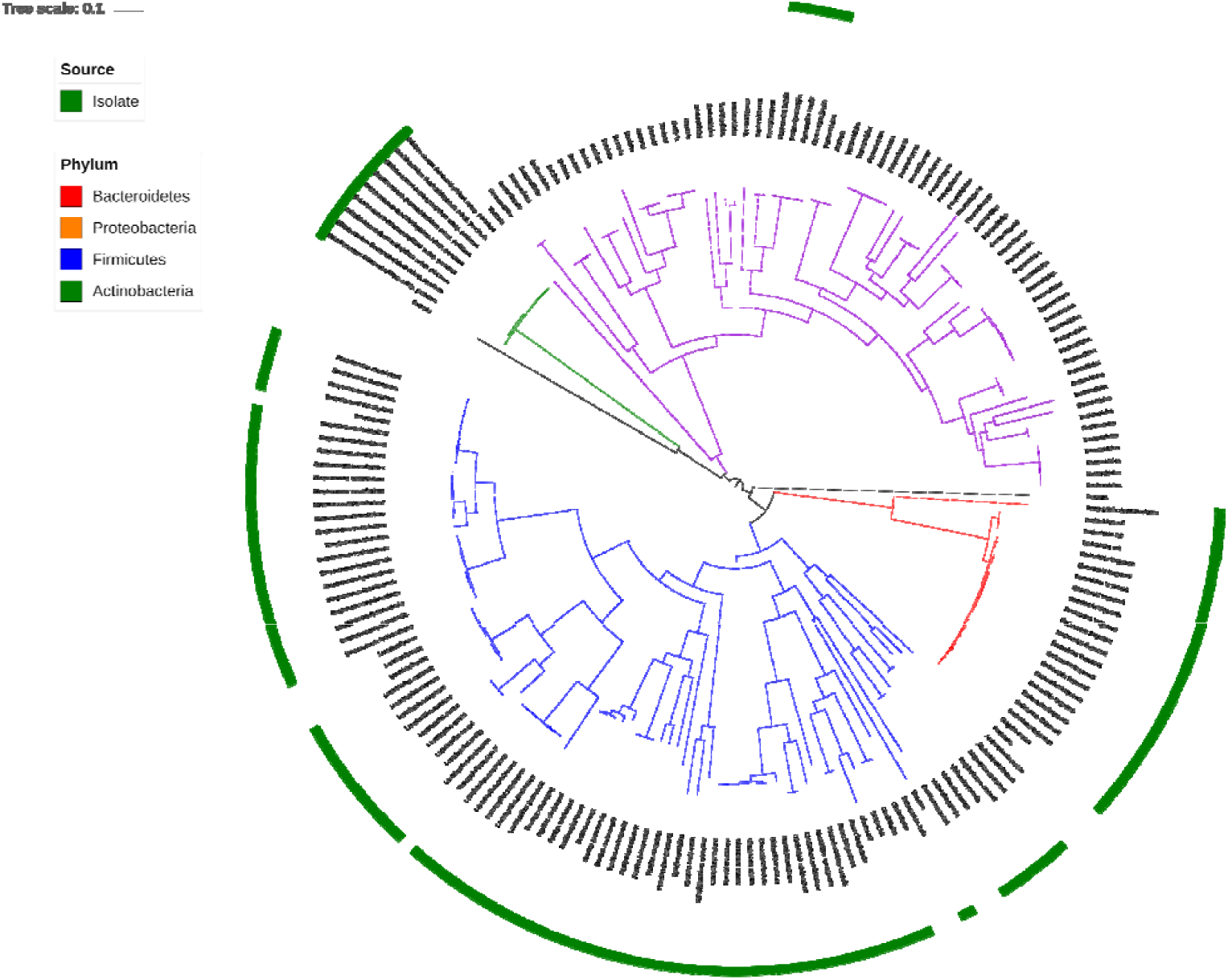
Maximum-likelihood tree of metagenomic bins and culture genomes. Tree was constructed from a nucleotide alignment of 92 single-marker genes. General time reversible (GTR) was chosen as the substitution model in tree construction.

**Fig 6.**
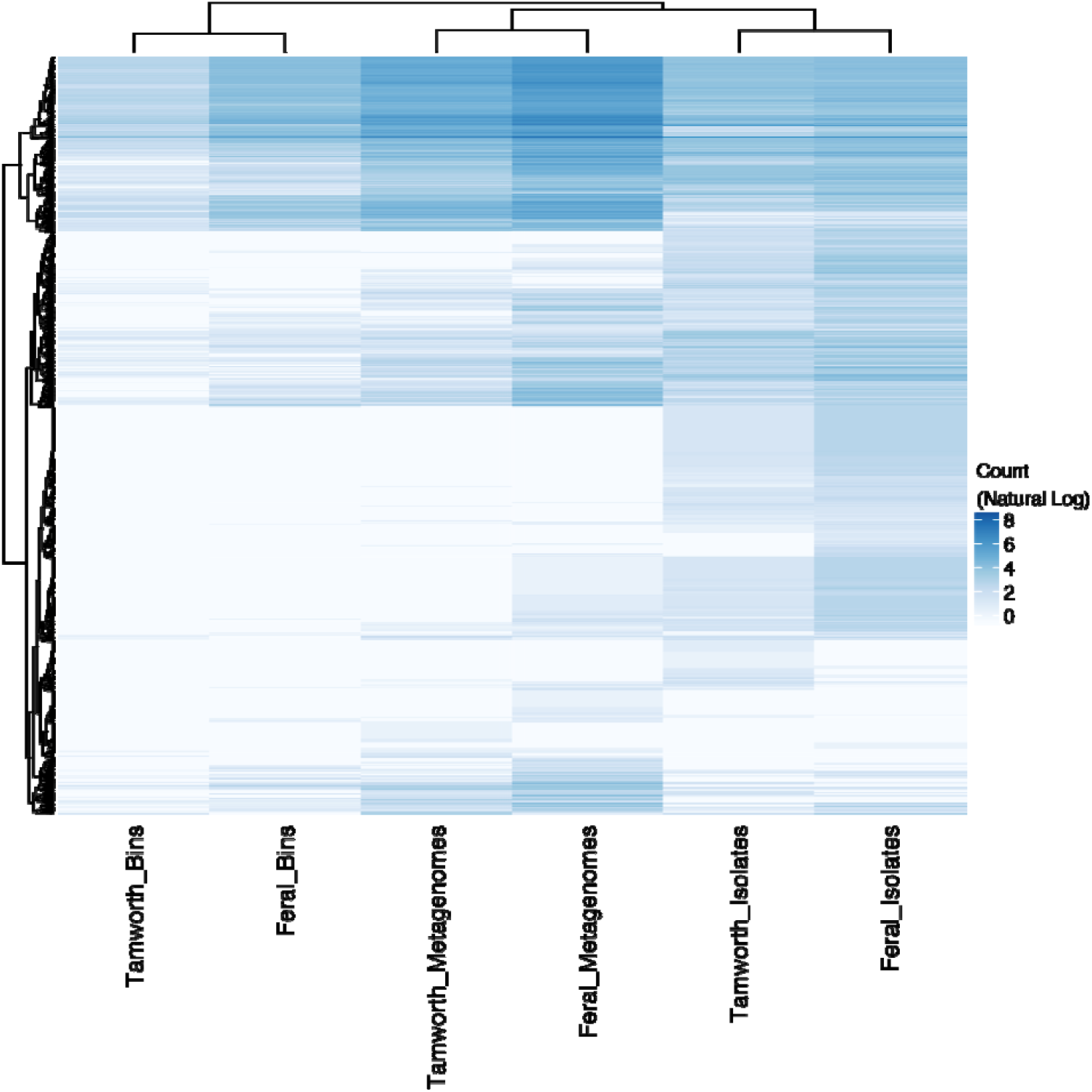
KEGG annotation of open frames from metagenomic assemblies, bins, and culture genomes. Rows and columns are clustered using an average linkage method. KEGG annotations counts are represented as the natural log to increase the clarity of the figure.

## Discussion

Metagenomic and culture analysis revealed that despite the “organic” raising of the Tamworth pigs studied (ability to forage, no antimicrobials), their microbiome does not resemble that of feral pigs. While it is true that the two most abundant genera were the same for both sources, *Prevotella* and *Bacteroides*, the Tamworth pigs examined were rather homogenous in microbiome composition and were dominated by the genus *Prevotella*. An increase of *Prevotella* in human samples has been attributed to increase of dietary fiber (David, et al. 2014: 559-63, De Filippo, et al. 2010: 14691-6, Smith, et al. 2013: 548-54). It was noted that the Tamworth pigs were fed a high forage diet and were bedded on alfalfa straw. The high dietary fiber intake in Tamworth pigs may be responsible for the high levels of Prevotella and could account for the lower diversity values compared to feral pigs. The exact diet of the feral pigs is unknown, but it has been observed that a major portion of feral pigs diet in Texas is composed of vegetation (Taylor and Hellgren 1997: 33-9). Prevotella has been identified as the most abundant genus in the swine microbiome to date (Holman, et al. 2017: e00004-17) and the sample size is simply too small to discern whether the large dominance of *Prevotella* in Tamworth pigs is breed or diet specific in nature.

Despite the high abundance of *Prevotella* in both Tamworth and feral pigs, and being the most abundant genus in pigs, the sampling strategy we employed only isolated seven *Prevotella* isolates from Tamworth samples (7/500, 1.4%) and no *Prevotella* was isolated from the feral inoculum. In contrast, several genera including *Lactobacillus, Escherchia, Streptococcous*, and *Bifidobacterium* were overrepresented in culture samples compared to metagenomic sequencing. Our culture results align with an early culture examination of the pig microbiome where the two most abundant isolates cultured were gram-positive cocci and *Lactobacillus* (Russell 1978: 187-93). Both our work and the earlier work relied upon complex media derived largely of peptone digests. As Prevotella is associated with an increase of dietary fiber, work will be needed to develop a defined media that is not based upon peptides such as yBHI. Culturomic techniques have largely focused on human fecal samples. Such studies have been wildly successful in culturing many bacteria that were previously thought to be “unculturable” (Browne, et al. 2016: 543, Lagier, et al. 2016: 16203). Many of the techniques rely upon anaerobic plating onto multiple media formulations, selection of single colonies, and identification. While the multiple media approach generates a higher number of taxa, one study isolated over 1,300 species (Lagier, et al. 2016: 16203), creating multiple media formulations can be expensive and time-consuming. Additionally, bacteria isolated from different media may not grow together on a common media, forfeiting any combined in vitro experimentation. Given the importance of swine in global agricultural, coordinated culture efforts are needed to develop defined community models. Such reduced communities will help to uncover the impact of major ecological principles (drift, selection, speciation, dispersion) at work in the swine hindgut.

One of the original motivations for this work was to establish the microbiotas of pigs outside of traditional agricultural processes. Remarkably, it is the Tamworth pigs and not the feral pigs that depart from the pig microbiota previously established (Holman, et al. 2017: e00004-17). The ratio of Firmicutes to Bacteroidetes is roughly equal in feral samples and that result aligns with agricultural animals. Tamworth at the phylum level is dominated by Bacteroidetes. Turning to the genus level, the top genus from both sources, *Prevotella*, aligns with the most abundant genus isolated from agricultural animals (Holman, et al. 2017: e00004-17). The genus *Bacteroides* is the second most abundant genus identified in both Tamworth and feral samples and is found in greater abundance than in conventionally-reared agricultural animals. The nearly identical distribution of *Bacteroides* between Tamworth and feral, and the discrepancy between conventionally-reared agricultural animals may indicate that traditional agricultural processes are negatively selecting for the genus. It has been shown that after weening *Bacteroides* levels plummet in growing pigs and are supplanted by *Prevotella* (Frese, et al. 2015: 28-). Yet in our samples a stable population of *Bacteroides* has persisted. It should be noted that the discrepancy may be accounted for by differing identification methods, metagenomics vs amplicon sequencing, or could be an artifact of sampling size.

The Tamworth pigs harbored more AMR homologues than the feral pigs despite no antimicrobials being provide. It has been shown previously that organically raised pigs harbor significantly more chlortetracycline resistant isolates than feral pigs (Stanton, et al. 2011: 7167). The previous report and our findings indicate that feral pigs are not a significant reservoir of AMR genes. However, the presence of AMR genes in Tamworth pigs may be contributed to recombination. Previous work has established that AMR genes may cluster together with mobile genetic elements and that pigs typically harbor genes conferring resistance to agents not typically used on a particular farm (Johnson, et al. 2016: e02214-15).

Recent studies have proposed metagenomic binning as a culture-independent method to extract genomes from samples (Albertsen, et al. 2013: 533, Pasolli, et al. 2019: 649-62.e20, Tully, et al. 2018: 170203, Wang, et al. 2019: 48). However, one of the main pitfalls of metagenomic binning is that metagenomic assemblers struggle to assemble contigs of closely related taxa, especially if the organisms are found in low abundance (Ayling, et al. 2019). With knowledge now that strain-level variation occurs in species of the microbiome (Lloyd-Price, et al. 2017: 61-6), targeted culture efforts are needed to confirm that strain variation observed in metagenomic data is not simply due to assembler bias. Also, a large portion of genes were not annotated in metagenomic assemblies that were identified in culture isolates. We propose a wholistic approach where metagenomic sequencing coupled with high-throughput culture strategies can effectively cover the shortcomings of either technique, leading to a more complete method of microbiome sampling.

## Acknowledgements

Computations supporting this project were performed on High-Performance Computing systems managed by Research Computing Group, part of the Division of Technology and Security at South Dakota State University. This work was in part supported by the grants from the South Dakota Governors Office of Economic Development (SD-GOED) and the United States Department of Agriculture (grant numbers SD00H532-14 and SD00R646-18) awarded to JS. We greatfully thank Tommy Harrison, Jeff Hopper and Brett Grogan for their help in collecting feral pig gut microbiota samples. We are saddened by the untimely passing away of Tomas Harrison during the course of this study. We dedicate this manuscript to Tom’s memory.

